# Sapropterin Treatment Prevents Congenital Heart Defects Induced by Pregestational Diabetes in Mice

**DOI:** 10.1101/304006

**Authors:** Anish Engineer, Tana Saiyin, Xiangru Lu, Andrew S. Kucey, Brad L. Urquhart, Thomas A. Drysdale, Kambiz Norozi, Qingping Feng

## Abstract

**Aims:** Tetrahydrobiopterin (BH4) is a co-factor of endothelial nitric oxide synthase (eNOS), which is critical to embryonic heart development. We aimed to study the effects of sapropterin (Kuvan®), an orally active synthetic form of BH4 on eNOS uncoupling and congenital heart defects (CHDs) induced by pregestational diabetes in mice.

**Methods:** Adult female mice were induced to pregestational diabetes by streptozotocin and bred with normal males to produce offspring. Pregnant mice were treated with sapropterin or vehicle during gestation. CHDs were identified by histological analysis. Cell proliferation, eNOS dimerization and reactive oxygen species (ROS) production were assessed in the fetal heart.

**Results:** Pregestational diabetes results in a spectrum of CHDs in their offspring. Oral treatment with sapropterin in the diabetic dams significantly decreased the incidence of CHDs from 59% to 27% and major abnormalities, such as atrioventricular septal defect and double outlet right ventricle were absent in the sapropterin treated group. Lineage tracing reveals that pregestational diabetes results in decreased commitment of second heart field progenitors to the outflow tract, endocardial cushions, and ventricular myocardium of the fetal heart. Notably, decreased cell proliferation and cardiac transcription factor expression induced by maternal diabetes were normalized with sapropterin treatment. Furthermore, sapropterin administration in the diabetic dams increased eNOS dimerization and lowered ROS levels in the fetal heart.

**Conclusions:** Sapropterin treatment in the diabetic mothers improves eNOS coupling, increases cell proliferation and prevents the development of CHDs in the offspring. Thus, sapropterin may have therapeutic potential in preventing CHDs in pregestational diabetes.

## INTRODUCTION

Congenital heart defects (CHDs) are the most common structural birth defect, occurring in 1-5% of live births, making them the leading cause of death in the first year of infant life [1, 2]. The prevalence of CHDs has been rapidly increasing [3], and it is estimated that approximately 2.4 million Americans, including 1 million children, are living with a congenital malformation of the heart [2]. CHDs are formed when complex cellular and molecular processes underlying embryonic heart development are disturbed. The heart is developed from three pools of progenitor cells: the first heart field (FHF), the second heart field (SHF) and cardiac neural crest (CNC) [4]. The FHF progenitors initially form the primary heart tube. SHF cells are then added to the heart tube to form the right ventricle, give rise to myocardial and endothelial cells of the outflow tract and semilunar valves, as well as the vascular smooth muscle cells at the base of the aorta and pulmonary trunk. The CNC cells contribute to septation of the outflow tract and remodeling of semilunar valves, while the left ventricle is mainly formed from FHF progenitors. The SHF is particularly significant to CHDs as many common cardiac abnormalities including atrial and ventricular septal defects, cardiac valve malformation, double outlet right ventricle and truncus arteriosus, are caused by defects in SHF progenitors [4].

Pregestational diabetes (type 1 or 2) in the mother increases the risk of a CHD in the child by four-fold [5]. While good glycemic control in diabetic mothers lowers the risk, the incident of CHDs in their children is still higher than the general population [6, 7]. The prevalence of pregestational diabetes has nearly doubled from 0.58% to 1.06% from 1996 to 2014 [8]. As the prevalence of pregestational diabetes further increases in women during their reproductive age, more individuals will be born with CHDs, inevitably placing a large burden on the healthcare system [9].

Uncontrolled maternal diabetes is not conducive to proper gestation. Hyperglycemia leads to cellular oxidative stress through numerous pathways [10], which include increased electron transport chain flow resulting in mitochondrial dysfunction, non-enzymatic protein glycosylation and glucose auto-oxidation all contributing to reactive oxygen species (ROS) generation [11, 12]. Increased oxidative stress can lead to the inactivation of many molecules and proteins necessary for proper heart development. Endothelial nitric oxide synthase (eNOS) is intimately regulated by redox balance within the cell and is vital for cardiogenesis [13]. eNOS expression in the embryonic heart regulates cell growth and protects early cardiac progenitors against apoptosis [14]. The importance of eNOS in heart development has been demonstrated in eNOS^-/-^ mice by a spectrum of cardiovascular anomalies such as ventricular septal defects (VSDs), valvular malformations and hypoplastic coronary arteries [14-16].

Tetrahydrobiopterin (BH4) has antioxidant properties and is a critical co-factor for eNOS function [17]. It is required for eNOS dimer stabilization and is an allosteric modulator of arginine binding to the active site [18]. In states of oxidative stress, BH4 levels decline, and eNOS is uncoupled, resulting in decreased NO synthesis and increased superoxide production, perpetuating the oxidative environment of the cell [19]. The production of ROS is amplified by this feedback loop, further inducing eNOS uncoupling. Treatment with BH4 has been shown to recouple eNOS and improve vascular endothelial function in diabetes [20, 21]. However, the potential of BH4 to reduce the severity and incidence of CHDs is not known. Sapropterin dihydrochloride (Kuvan®) is an orally active synthetic form of BH4 and an FDA approved drug for the treatment of phenylketonuria [22]. The present study was aimed to examine the effects of sapropterin in mice with pregestational diabetes. We hypothesized that sapropterin treatment during gestation re-couples eNOS, improves cell proliferation in SHF derived cells, and reduces CHD incidence in the offspring of mice with pregestational diabetes.

## METHODS

### Animals

All procedures were performed in accordance with the Canadian Council on Animal Care guidelines and approved by the Animal Care Committee at Western University. C57BL/6 wild type and *Rosa26^mTmG^* mice were purchased from Jackson Laboratory (Bar Harbor, Maine). *Mef2c^cre/+^* embryos were obtained from Mutant Mouse Regional Resource Center (Chapel Hill, NC) and rederived. All animals were housed in a 12-hour light/dark cycle and given ad libitum access to standard chow and water. A breeding program was established to generate embryonic, fetal and post-natal mice.

### Induction of Diabetes and Sapropterin Treatment

Female C57BL/6 mice, 8 to 10 weeks old were made diabetic through five consecutive daily injections of streptozotocin (STZ, 50 mg/kg body weight, IP, Sigma) freshly dissolved in sterile saline. Mice were randomly assigned to STZ (n=37) or saline treatment (n=19) groups. One week following the last STZ injection, non-fasting blood glucose levels were measured with a tail snip procedure using a glucose meter (One Touch Ultra2, LifeScan, Burnaby, BC). Mice were categorized as diabetic if blood glucose measurements exceeded 11 mmol/L, and were subsequently bred to 10 to 12 week old C57BL/6 male mice. In the morning when a vaginal plug was observed indicating embryonic day 0.5 (E0.5), the female diabetic mouse was placed in a separate cage with littermates. A cohort of diabetic and control female mice were treated with sapropterin dihydrochloride (Kuvan®, BioMarin Pharmaceutical Inc.) at a dose of 10 mg/kg body weight per day during gestation. Sapropterin was dissolved in sterile water and mixed with a small amount peanut butter, ensuring it was fully consumed by the mouse. Non-fasting blood glucose levels were monitored throughout pregnancy. To prevent hyperglycemia, a long acting form of insulin (Lantus®, Sanofi Aventis) was administered subcutaneously to a cohort diabetic dams (n=3) at a dose of 0.5 units/day.

### Histological and Immunohistochemical Analysis

Fetal samples were harvested at E10.5, E12.5 and E18.5 for histological and immunohistochemical analysis. To diagnose CHDs in E18.5 hearts, fetuses were decapitated and the isolated thorax was fixed overnight in 4% paraformaldehyde, dehydrated in ethanol and paraffin embedded. Samples were sectioned in 5 μm slices and hematoxylin/eosin (H&E) or toluidine blue stained to visualize morphology. Images were taken and analyzed using a light microscope (Observer D1, Zeiss, Germany). Embryonic samples at E10.5 and E12.5 were fixed in 4% paraformaldehyde for 1 and 2 hours, respectively, and processed as described above. Immunostaining to analyze cell proliferation at E10.5 using anti-phosphohistone H3 (pHH3) antibody (1:1000, Abcam), and sex determination at E18.5 using anti-sex-determining region Y protein antibody (1:200, Santa Cruz) were performed after antigen retrieval in citrate buffer (10 mmol/L, pH 6). This was followed by incubation with biotinylated goat anti-mouse IgG (1:300, Vector Laboratories) secondary antibody. The signal was amplified by the ABC reagent (Vector Laboratories) allowing for visualization through 3-3’ di-aminobenzidine tetrahydrochloride (DAB, Sigma) with hematoxylin as a counterstain. Blinded pHH3^+^ cell counts within the outflow tract (OFT) were taken from at least 3 heart sections per heart and normalized to OFT length.

### Lineage Tracing the Second Heart Field (SHF)

Fate mapping of SHF progenitors was performed using the SHF specific *Mef2c^cre/+^* transgenic mouse and the global double fluorescent *Cre* reporter line *Rosa26^mTmG^* that has *LoxP* sites on either side of a tomato-red fluorescence membrane protein (mT) cassette, which is proceeded by a green fluorescence protein (GFP, mG) cassette. In the *Mef2c^Cre/+^* transgenic line with C57BL/6 background, elements of the *Mef2c* promoter drives *Cre* recombinase expression in all SHF derived cells. When crossed with the *Rosa26^mTmG^* mice this results in a GFP signal that is detected in all SHF derived cells [23, 24]. In all other tissues, the absence of the *Cre* results in mT expression and red fluorescence. Diabetes was induced in homozygous *Rosa26^mTmG^* females (8 to 10 weeks old) by STZ as described above. Hyperglycemic *Rosa26^mTmG^* female mice were crossed with *Mef2c^cre/+^;Rosa26^mTmG^* males to generate E9.5 and E12.5 *Mef2c^cre/+^;Rosa26^mTmG^* embryos, which were fixed, embedded and sectioned in the same manner as above. Immunostaining for membrane bound GFP was conducted using an anti-GFP (1:500, Abcam) primary antibody, followed by biotinylated goat anti-rabbit IgG (1:300, Vector Laboratories) secondary antibody using DAB for visualization. The SHF derived cells, which are GFP^+^, were blindly quantified and compared between control and diabetic groups.

### Analysis of Superoxide Levels

Hearts collected form E12.5 fetuses from all four groups were cryo-sectioned (CM1950, Leica, Germany) into 8 μm thick slices and placed onto slides. Samples were incubated with 2 μM dihydroethidium (DHE) probe (Invitrogen Life Technologies, Burlington, Canada) for 30 minutes in a dark humidity chamber at 37 °C. After cover glass was mounted, DHE fluorescence signals were visualized using a fluorescence microscope (Observer D1, Zeiss, Germany). Five – 8 images from each sample were captured at fixed exposure times for all groups and the fluorescence intensity per myocardial area was blindly quantified using AxioVision software.

### Measurement of Cardiac Function

Pregnant dams were anesthetized and M-mode echocardiography images of E18.5 embryonic hearts were recorded using the Vevo 2100 ultrasound imaging system with a MS 700 transducer (VisualSonics, Toronto, Canada) as previously described [15, 16]. The end diastolic left ventricular internal diameter and end systolic left ventricular internal diameter were measured from the short-axis M-mode images to calculate LV ejection fraction and fractional shortening.

### Western Blotting for eNOS Dimers and Monomers

Ventricular myocardial tissues from E12.5 hearts were dissected in PBS and used for measurement of eNOS dimerization. In order to not disrupt the dimer, proteins were isolated from three pooled hearts from each group via sonication using a non-reducing lysis buffer, on ice [25]. Protein lysates without boiling were run on an 8% SDS-PAGE at 4 °C followed by transferring to a nitrocellulose membrane and immunoblotted with anti-eNOS polyclonal antibody (1:3000, Santa Cruz). This technique resulted in two distinct bands at 260 and 130 kDa, representing the eNOS dimer and monomer, respectively [25]. Proteins isolated from cultured coronary artery microvascular endothelial cells was boiled and separated with the sample, acting as an eNOS monomer size control.

### Real-time RT-PCR

Total RNA was isolated from E12.5 hearts using TRIzol reagent (Invitrogen). 200 nanograms of RNA were synthesized into cDNA with Moloney murine leukemia virus reverse transcriptase and random primers. Real-time PCR was conducted on cDNA using Evagreen qPCR MasterMix (Applied Biological Systems, Vancouver, BC). Primers were designed for Gata4, Gata5, Nkx2.5, Tbx5, GCH1, DHFR using the Primer3 software v 4.1.0 (Supplementary Table 1). Eppendorf Realplex (Eppendorf, Hamburg) was used to amplify samples for 35 cycles. Values were normalized to 28S ribosomal RNA, and mRNA levels were extrapolated through a comparative Ct method [16].

### Determination of Biopterin Levels

Tissue and plasma biopterin levels were determined using ultra-performance liquid chromatography (UPLC) coupled to mass spectrometry. Briefly, E12.5 whole embryos were harvested and blood samples were collected from the dams. Plasma samples were then diluted 1:4 with 150 μl of acetonitrile containing internal standard (100 μM aminopimilic acid and 2.5 μM chlorpromazine) before injection into the ultra-performance liquid chromatography (UPLC) instrument. Total BH4 and BH2 levels were assessed using a Waters Acuity I Class UPLC system coupled to a XEVO G2-S quadrupole time-of-flight mass spectrometer (Waters Corporation: Milford, Massachusetts) as previously described [26].

### Statistical Analysis

Data are presented as the mean ± SEM. Statistical analysis was performed using GraphPad Prism (Version 5, GraphPad Software, La Jolla, CA, USA). Multiple group comparisons was conducted using two-way analysis of variance (ANOVA) followed by the Bonferroni post-hoc test. Two groups were compared using unpaired Student’s t-test. The incidence of congenital heart malformations was assessed with Chi-square test. Differences were deemed significant with *P*<0.05.

## RESULTS

### Effects of sapropterin on blood glucose, fertility, litter size and biopterin levels in pregestational diabetes

This study was conducted in the same pregestational diabetes model we recently employed [27, 28]. One week following the final administration of STZ, female mice with random blood glucose levels >11 mmol/L were bred with normal adult males. During gestation, blood glucose levels in the diabetic dams were progressively increased from E0.5-18.5 compared to both sapropterin treated and untreated control mice (Fig. 1A). Treatment with insulin but not sapropterin in the diabetic mice restored blood glucose to normal levels. Diabetic dams had a significant lower fertility rate (38%) compared to controls (82%, *P*<0.05), which was improved to 64% by sapropterin treatment (Fig. 1B).

**Figure 1.**
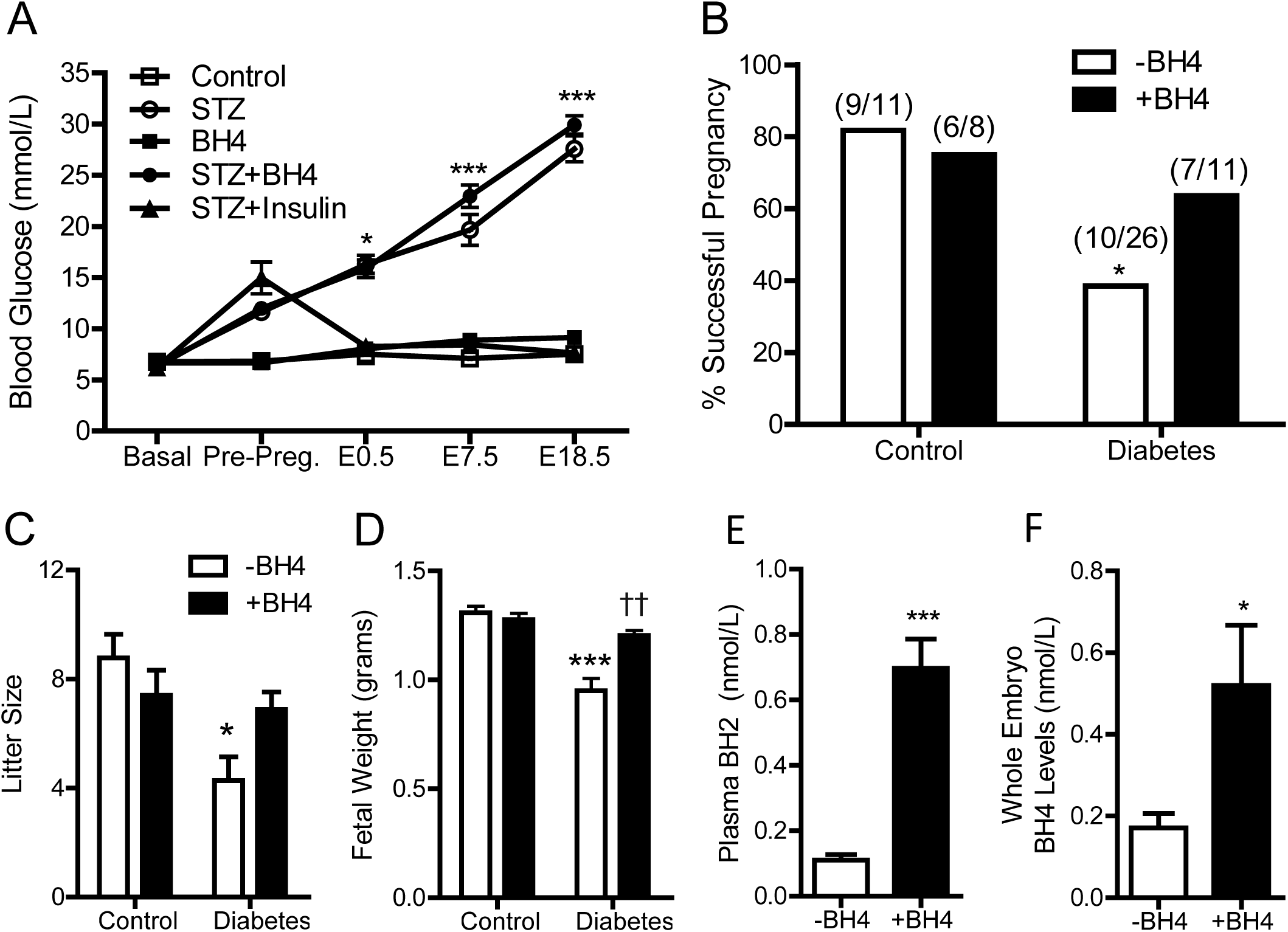
Blood glucose levels of pregnant mice, percent successful plugs, litter size, fetal body weight and biopterin levels. (A) Non-fasting blood glucose levels from before mating (basal) to E18.5 during pregnancy in STZ-treated and control female mice with and without sapropterin (BH4) administration (n = 4-8 mice per group). (B) The percent of successful pregnancy. The numbers in the brackets indicate the number of successful pregnancy at the day of fetus harvest to total plugs. (C) The offspring litter size measured at the day of fetus harvest (n = 5-11 litters per group). (D) Fetal body weight at E18.5 (n = 5-8 fetuses per group). (E-F) Levels of BH2 in maternal plasma, and BH4 in diabetic embryos with or without BH4 administration (n = 3-9 mice per group). \**P*<0.05; \*\*\**P*<0.001 vs. corresponding controls, and ††*P*<0.001 vs. untreated diabetes. Data are means ± SEM in A, C-F.

Diabetic dams had a significantly smaller litter size (*P* < 0.01), which was improved by sapropterin treatment (Fig. 1C). Indeed, absorbed or dead fetuses were more commonly seen in utero in diabetic dams. Fetal body weight was significantly lower from diabetic dams than controls (*P*<0.001), and was restored to normal weight with sapropterin (*P*<0.001, Fig. 1D). To assess biopterin levels following oral sapropterin administration, maternal blood and embryos were collected at E12.5. Our data show that sapropterin treatment significantly increased plasma BH2 levels in the diabetic mothers (*P*<0.01, Fig. 1E) and BH4 levels in E12.5 embryos of diabetic dams (Fig. 1F).

### Sapropterin prevents CHDs induced by pregestational diabetes

CHDs were observed in 59.4% of offspring from diabetic dams (Table 1 and Fig. 2). Normal control heart sections are shown in Fig. 2A-C. Septal defects constituted a large proportion of the anomalies with 47.4% atrial septal defects (ASD, Fig. 2D) and 37.8% ventricular septal defects (VSD, Fig. 2E). Hypoplastic left heart was seen in 21.1% of the diabetic offspring. One fetus had an atrioventricular septal defect (AVSD). Outflow tract (OFT) defects were also common in offspring from diabetic mothers. A double outlet right ventricle (DORV) was present in one of the offspring (Fig. 2F) and truncus arteriosus was seen in another fetus (Fig. 2G). Maternal diabetes also resulted in cardiac valve defects with 37.8% thickened aortic valves (Fig. 2H) and 29.7% thickened pulmonary valves (Fig. 2I). Finally, 24.3% of hearts from diabetic mothers displayed a narrowing aorta (Fig. 2H). Treatment with sapropterin to diabetic dams during pregnancy significantly decreased the overall incidence of CHDs to 26.5% (Table 1). Specifically, offspring from sapropterin-treated diabetic mothers show a significantly lower incidence of ASD and thickened aortic valves, and did not display any VSD, AVSD, truncus arteriosus, DORV or hypoplastic left heart (Table 1, Fig. 2J-O). No CHDs were seen in diabetic dams who received a daily dose of insulin to maintain normal blood glucose levels, indicating that CHDs seen in the diabetic offspring were induced through hyperglycemia (Table 1, Fig. 2P-R). The incidence of CHDs was significantly higher in males than females (65% vs. 47%, P<0.05). Furthermore, sapropterin treatment was more effective in reducing the incidence of CHDs to 16% in females compared to 35% in males (*P*<0.05, Fig. 2S). Cardiac function of E18.5 fetuses was assessed through echocardiography. LV fractional shortening and ejection fraction were significantly decreased in offspring of diabetic dams, and were recovered by sapropterin treatment (*P*<0.001, Fig. 2T and U). Pregestational diabetes also induced neural tube defects (NTD) in our model. A case of exencephaly in the offspring of mother with pregestational diabetes is shown in supplementary Figure 1.

**Table 1.** The rate of congenital heart defects in the offspring of diabetic and control mothers with and without sapropterin (BH4) treatment. Data was analyzed using the Chi-square test. \**P* <0.05, \*\**P* <0.001 vs. untreated control, †*P* <0.05, ††*P* <0.001 vs. untreated diabetes. ASD: atrial septal defect, VSD: ventricular septal defect, AVSD: atrioventricular septal defect, DORV: double outlet right ventricle. N indicates total number of fetuses, dams are litter numbers.

|  | Control |  | STZ |  | BH4 |  | STZ+BH4 |  | STZ+Insulin |  |
| --- | --- | --- | --- | --- | --- | --- | --- | --- | --- | --- |
|  | N | dams | N | dams | N | dams | N | dams | N | dams |
|  | 14 | 3 | 38 | 9 | 23 | 4 | 34 | 5 | 20 | 3 |
|  | n | % | n | % | n | % | n | % | n | % |
| Normal | 14 | 100 | 16 | 40.6** | 23 | 100 | 25 | 73.5† | 20 | 100 |
| Abnormal | 0 | 0 | 22 | 59.4** | 0 | 0 | 9 | 26.5†† | 0 | 0 |
| ASD | 0 | 0 | 18 | 47.4** | 0 | 0 | 9 | 26.5† | 0 | 0 |
| VSD | 0 | 0 | 14 | 37.8** | 0 | 0 | 0 | 0.0†† | 0 | 0 |
| Truncus Arteriosus | 0 | 0 | 1 | 2.7 | 0 | 0 | 0 | 0.0 | 0 | 0 |
| AVSD | 0 | 0 | 2 | 5.4* | 0 | 0 | 0 | 0.0† | 0 | 0 |
| DORV | 0 | 0 | 1 | 2.7 | 0 | 0 | 0 | 0.0 | 0 | 0 |
| Pulmonary Valve Stenosis | 0 | 0 | 14 | 37.8** | 0 | 0 | 0 | 0.0†† | 0 | 0 |
| Aortic Valve Stenosis | 0 | 0 | 11 | 29.7** | 0 | 0 | 1 | 2.9†† | 0 | 0 |
| RV Hypertrophy | 0 | 0 | 9 | 24.3** | 0 | 0 | 0 | 0.0†† | 0 | 0 |
| Hypoplastic Left Heart | 0 | 0 | 8 | 21.1** | 0 | 0 | 0 | 0.0†† | 0 | 0 |
| Narrowing of the Aorta | 0 | 0 | 9 | 24.3** | 0 | 0 | 0 | 0.0†† | 0 | 0 |

**Figure 2.**
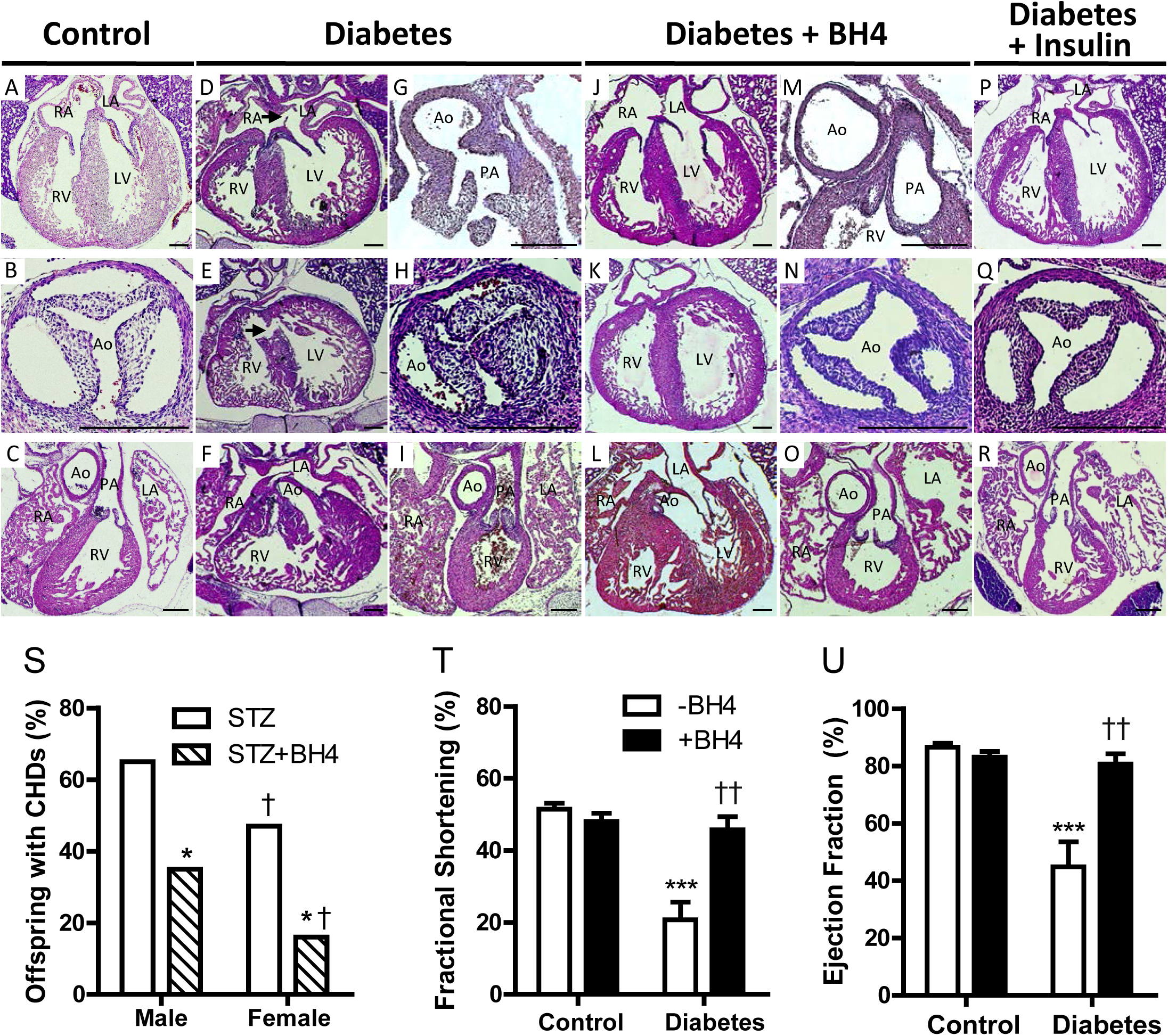
Effects of sapropterin (BH4) on congenital heart defects induced by pregestational diabetes. Representative histological sections of E18.5 hearts from offspring of diabetic mothers with and without BH4 treatment. (A-C) show hearts sections from controls. Pregestational diabetes resulted in atrial septal defect (D), ventricular septal defect (E), double outlet right ventricle (F), truncus arteriosus (G), thickened aortic (H) and pulmonary (I) valves. BH4 treatment in diabetic dams shows normal heart morphology (J-O). Diabetic dams on insulin therapy to control blood glucose levels show normal fetal heart morphology (P-R). Scale bars are 200 μm. (S) Incidence of CHDs as percent of offspring separated by sex at E18.5 from diabetic dams with or without BH4 treatment. \**P*<0.05 vs. STZ group of corresponding sex; †*P*<0.05 vs. corresponding males. (T-U) LV fractional shortening and ejection fraction assessed by echocardiography in E18.5 offspring (n = 5-6 fetuses per group). \*\*\**P*<0.001 vs. corresponding controls, and ††*P*<0.001 vs. untreated diabetes. Data are means ± SEM

### Sapropterin prevents myocardial and valvular abnormalities induced by pregestational diabetes

The free walls of the right and left ventricle at E18.5 were measured at the mid-ventricular region (Fig. 3A). The RV and LV myocardium was thinner in the offspring of diabetic mothers compared to those of control offspring, which was prevented by sapropterin treatment (*P*<0.001, Fig. 3B and C). Additionally, since valve leaflets were thickened (Fig. 2H and I), we assessed glycosaminoglycans, a component of extracellular matrix in the cardiac valves using toluidine blue staining. Our data show that glycosaminoglycans occupied a greater space in the leaflets of aortic and pulmonary valves (light purple color) from E18.5 fetal hearts of diabetic mothers compared to controls (Fig. 4A and B). Treatment with sapropterin decreased these extracellular polysaccharides in both aortic and pulmonary valves (*P*<0.01, Fig. 4D and E). There was no significant difference in glycosaminoglycan content in the mitral valve, however the distal tip of mitral valve was thicker in the offspring of diabetic dams (*P*<0.01, Fig. 4F). Notably, the area of aortic orifice and diameter of pulmonary artery were significantly smaller in hearts from diabetic mothers compared to controls (*P*<0.01, Fig. 4G and H).

**Figure 3.**
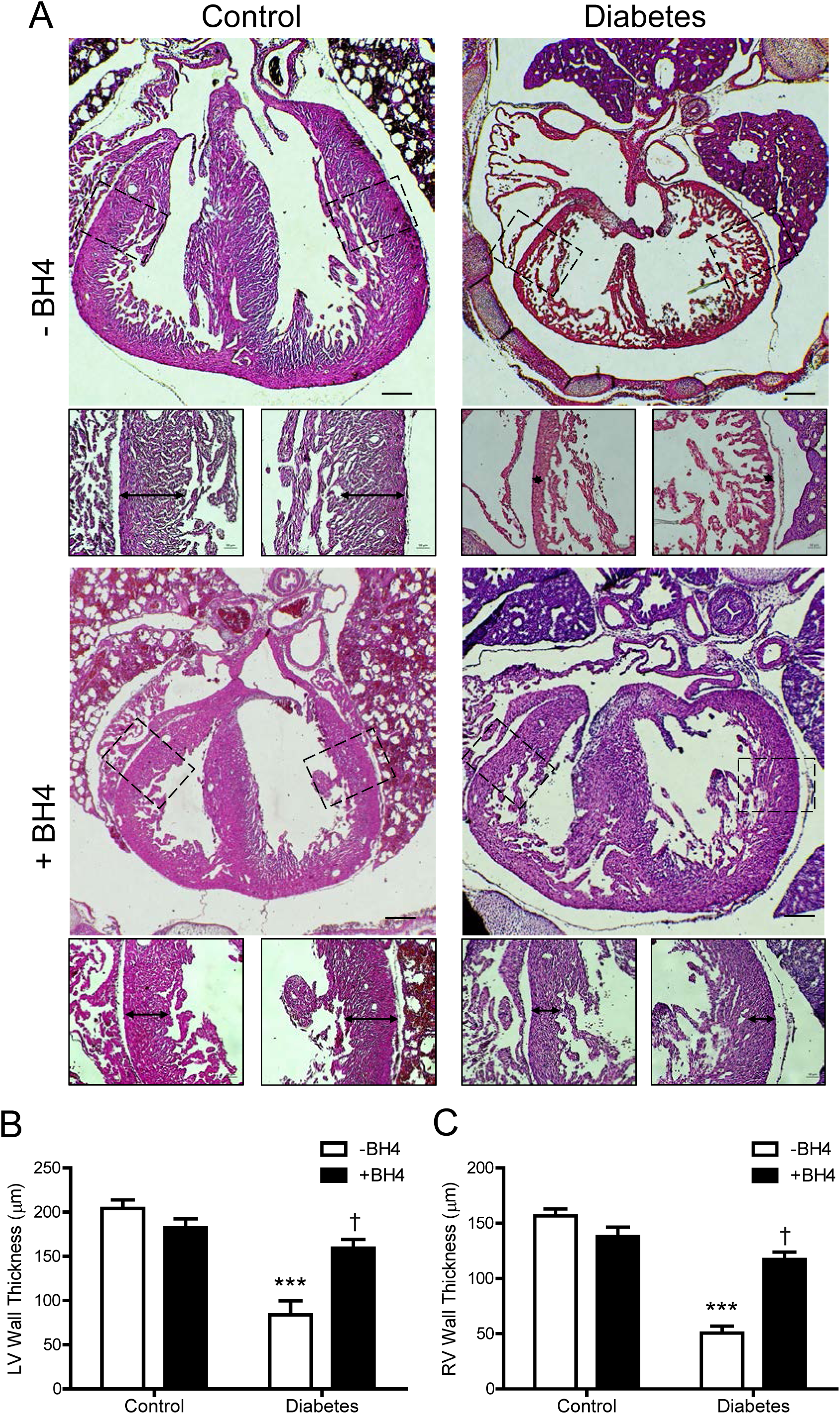
Effects of sapropterin (BH4) on myocardialization of fetal hearts of pregestational diabetes. (A) Representative images of myocardial wall thickness in control and pregestational diabetes with and without BH4 treatment. Arrows indicate compact myocardium boundary from which measurements were obtained. (B-C) Quantification of right and left ventricular myocardial thickness, respectively. *n* = 6 per group from 3 - 6 litters, \*\*\**P*<0.01 vs. untreated control, †*P*<0.01 vs. untreated diabetes. Scale bars are 200 μm.

### Effects of sapropterin on outflow tract length and cell proliferation in the fetal heart

Since offspring of diabetic mothers show OFT defects, we aimed to investigate changes in cell proliferation at a critical stage in OFT formation. At E10.5 sagittal sections of whole embryos reveal a shortened OFT in hearts from diabetic mothers compared to control (*P*<0.001, Fig. 5A), which was restored with sapropterin treatment (*P*<0.05, Fig. 5E). To further analyze this change, phosphorylated histone H3, a marker for the mitotic phase of cell division, was used to compare the levels of proliferating cells in the OFT at E10.5 using the same hearts (Fig. 5B-D). Embryos from mice with pregestational diabetes had a significantly lower number of proliferating cells in the OFT (*P*<0.01, Fig. 5F-G). Sapropterin treatment was able to prevent impaired cell proliferation in the OFT in hearts from diabetic dams (*P*<0.001, Fig. 5F-G). Additionally, the number of proliferating cells in the ventricular myocardium of these hearts was significantly decreased in fetal hearts of diabetic dams, which was restored to normal levels with sapropterin treatment (*P*<0.01, Fig. 5D and G).

**Figure 4.**
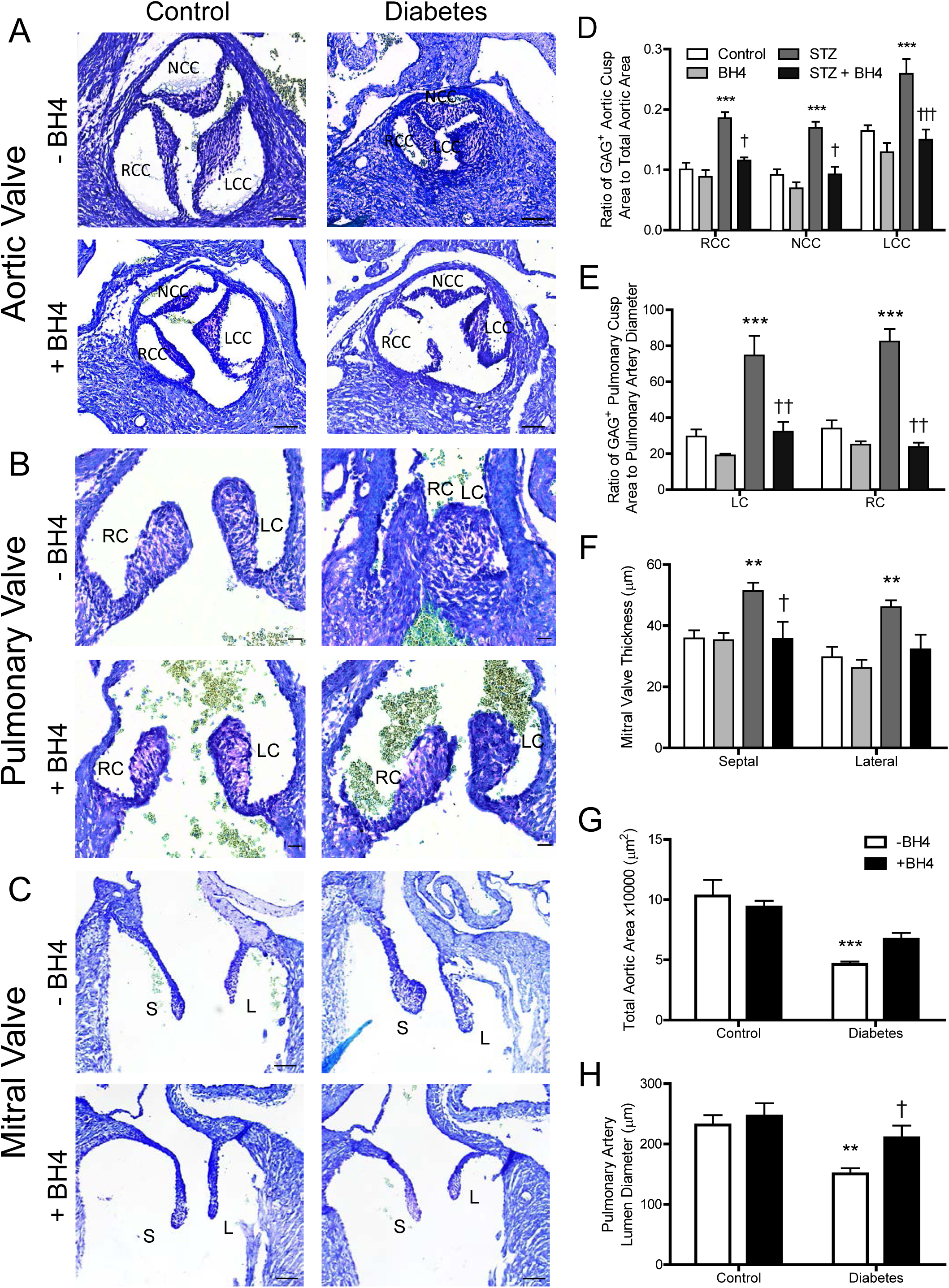
Effects of sapropterin (BH4) on aortic, pulmonary and mitral valve defects induced by pregestational diabetes. (A-C) Representative images of toluidine blue staining of glycosaminoglycans (GAG) in aortic, pulmonary, and mitral valves in E18.5 hearts. (D) The ratio of GAG+ area to total valve leaflet area. (E) Pulmonary valve leaflet thickness. (F) Mitral valve leaflet thickness. (G) Total aortic orifice area. (H) Pulmonary artery luminal diameter at the base of the orifice. LCC, left coronary cusp; NCC, noncoronary cusp; RCC, right coronary cusp; RC, right cusp; LC, left cusp; S, septal; L, lateral. Scale bars are 50, 20 and 20 μm in A, B and C, respectively. \**P*<0.05, \*\**P*<0.01 vs. controls. †*P*<0.01, ††*P*< 0.01 vs. untreated diabetes. n = 4-7 hearts per group from 3 - 4 litters.

**Figure 5.**
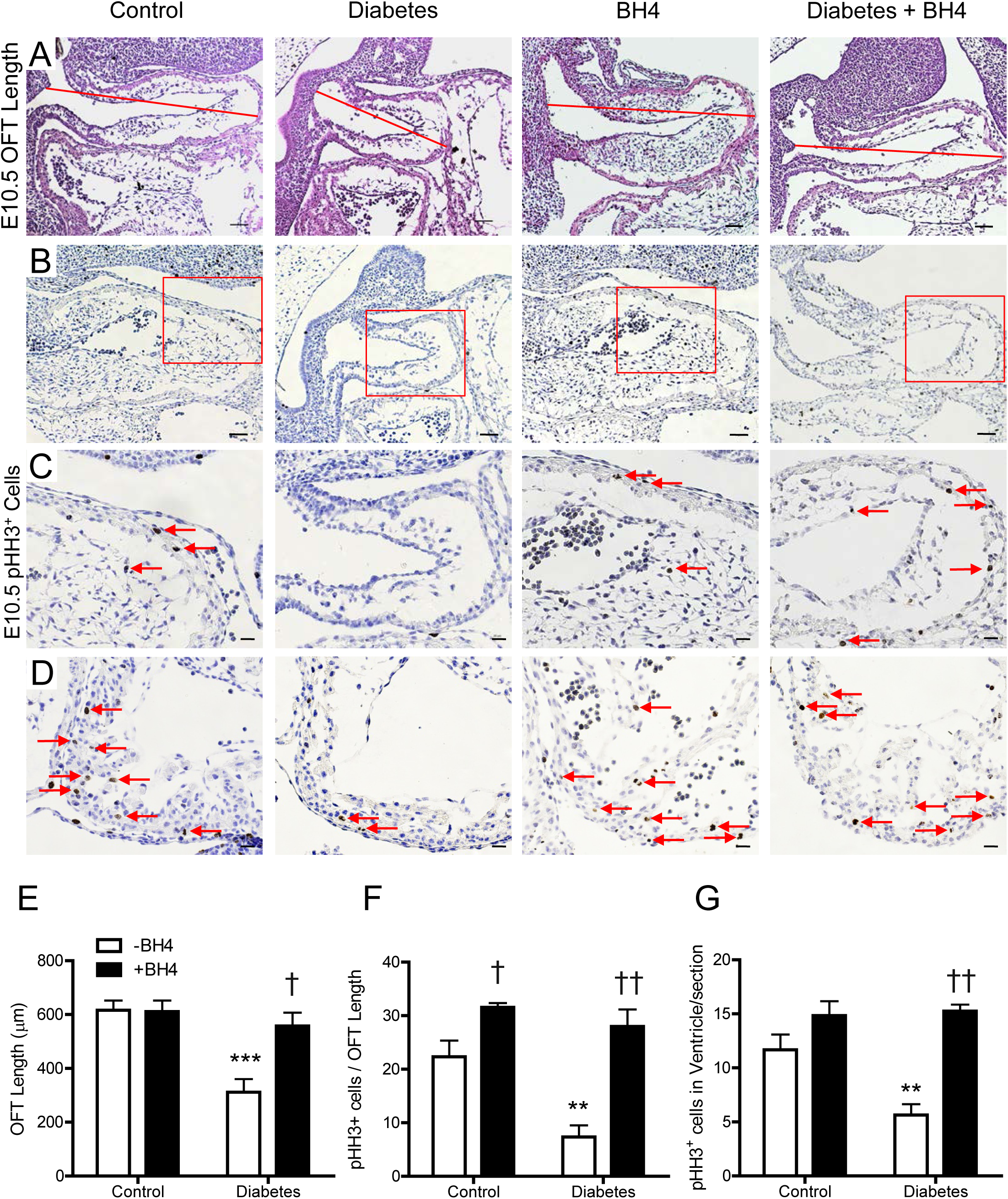
Effects of sapropterin (BH4) on outflow tract development and cell proliferation in E10.5 hearts of pregestational diabetes. (A) Length of the OFT was measured via sagittal sections according to the line. (B-D) Immunostaining for phosphorylated histone H3 marking proliferating cells in the OFT (B-C) and ventricular myocardium (D). Panel C is the magnification of boxed area in panel B. (E) Quantification of OFT length. (F) Quantification of pHH3^+^ cells in the OFT. (G) Quantification of pHH3^+^ cells in the ventricle. n = 3-6 hearts per group from 2 - 4 litters, \*\**P*<0.01 vs. untreated control, †*P*<0.05 vs. untreated control, ††*P*<0.01 vs. untreated diabetes. Scale bars are 50 μm and 20 μm, respectively.

### Fate-mapping of SHF derived cells in the fetal heart of diabetic mothers

In order to better understand the spectrum of malformations induced by maternal diabetes seen at birth, embryonic lineage tracing of the SHF was carried out. Fate mapping using *Mef2c-Cre* and the global double fluorescent Cre reporter line *Rosa26^mTmG^* identified all SHF derived cells as GFP^+^. At E9.5, the number of GFP^+^ SHF cells and total number of cells in the heart were significantly less than control (*P*<0.01, Fig. 6A, D and E). Furthermore, significantly less GFP^+^ SHF cells were infiltrated into the endocardial cushion at E12.5 in diabetic embryos compared to control (*P*<0.05, Fig. 6B andF). Additionally, E12.5 hearts from diabetic mothers had thinner ventricular walls with significantly less myocardial cell layers than controls (*P*<0.01, Fig. 6C and G).

**Figure 6.**
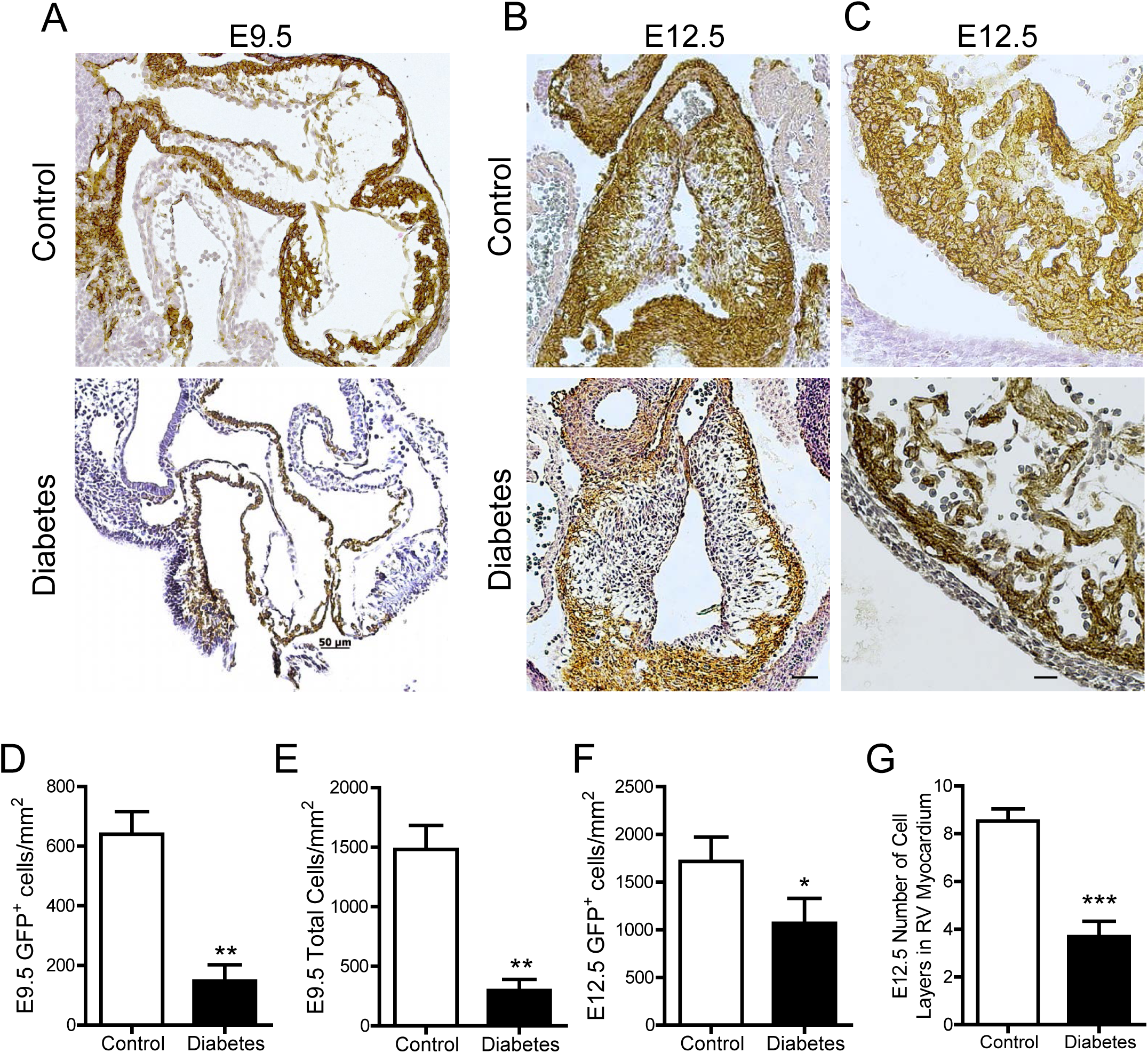
Effects of pregestational diabetes on SHF progenitor contribution to E9.5 and E12.5 hearts. Fate mapping using the *mT/mG* reporter showing GFP^+^ cells expressing Cre recombinase under the control of the anterior heart field specific Mef2c transcription factor. Representative sections of OFT of E9.5 (A) and E12.5 (B) hearts from control and diabetic mothers. (C) Representative sections showing the cell layers of the RV myocardium from E12.5 hearts. Quantification of SHF GFP^+^ cells (D) and total number of cells (E) in the OFT cushions at E9.5. Quantification of SHF GFP^+^ cells in the OFT cushions (F) and number of cell layers in the RV myocardium (G) at E12.5. N = 3 per group from 2 litters in D and E. N = 5 per group in F and G from 2 - 4 litters. \**P*<0.05, \*\**P*<0.01, \*\*\**P*<0.001 vs. control. Scale bars are 50 μm.

### Sapropterin prevents maternal diabetes-induced downregulation of regulators of heart development

Pregestational diabetes induces changes in gene expression in the developing heart [29]. To determine if key transcriptional regulators of heart development were altered under maternal diabetes and with sapropterin treatment, qPCR analysis was performed from E12.5 hearts. Our data show that the mRNA levels of Gata4, Tbx5, Nkx2.5, Gata5 and Bmp10 were significantly lower compared to controls (*P*<0.05, Fig. 7A-E). Treatment with sapropterin significantly improved mRNA levels of these transcription factors (*P*<0.001, Fig. 7A-E). Additionally, the expression of BH4 biosynthesis enzymes including GTP cyclohydrolase 1 (GCH1) and dihydrofolate reductase (DHFR) was significantly decreased in embryonic hearts from diabetic mothers (*P*<0.05, Fig. 7F and G). Of note, sapropterin treatment completely recovered the expression of DHFR (*P*<0.001, Fig. 7G).

**Figure 7.**
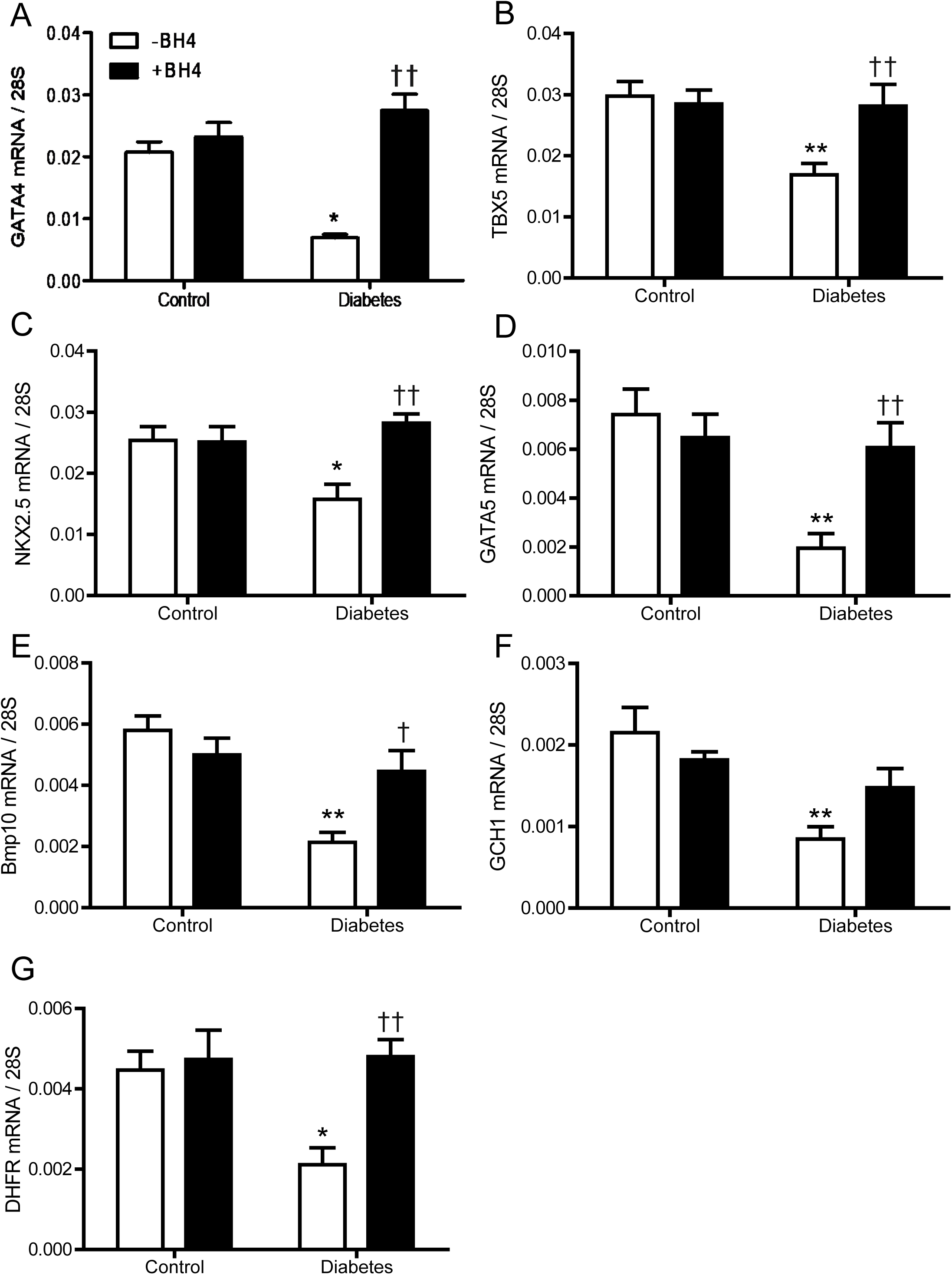
Effects of sapropterin (BH4) on molecular regulators of heart development. Real-time RT-PCR of cardiac transcription factors and regulators in E12.5 hearts of offspring from control and diabetic mothers. (A-E) The expression of *Gata4, Tbx5, Nkx2.5, Gata5* and *Bmp10* were significantly decreased under maternal diabetes and restored with BH4 treatment. (F-G) The expression of *GCH-1* and *DHFR* was significantly decreased with maternal diabetes. BH4 treatment rescues expression of *DHFR*. n = 4 - 7 hearts per group. \**P*<0.05, \*\**P*<0.01 vs. untreated control, and †*P*<0.05, ††*P*<0.01 vs. untreated diabetes.

### Sapropterin decreases oxidative stress and improves eNOS dimerization in fetal hearts of pregestational diabetes

To examine the effects of sapropterin on ROS in the developing heart, DHE probe was used to label the oxygen radical. Quantification of red fluorescence intensity indicates that superoxide generation was significantly elevated in embryonic hearts from mice with pregestational diabetes compared to control (*P*<0.05, Fig. 8A and B). Sapropterin treatment significantly reduced myocardial ROS levels to basal conditions (*P*<0.05 Fig. B). Finally, eNOS uncoupling has been implicated as a mechanism for endothelial dysfunction in diabetes [20]. To assess the effects of sapropterin on eNOS coupling, eNOS dimerization was analyzed using Western blotting in non-reducing conditions, which yields two bands at 260 and 130 kDa, representing the intact dimer and monomer of eNOS, respectively. Figure 8C shows a representative blot of the dimer and monomer of eNOS. In control conditions, there was a higher level of eNOS dimers than monomers, indicating a functional eNOS enzyme. However, embryonic hearts from diabetic dams show a significant decrease of dimer/monomer ratios compared to controls (*P*<0.05, Fig. 8D), which was restored by sapropterin treatment (*P*<0.01, Fig. 8D).

**Figure 8.**
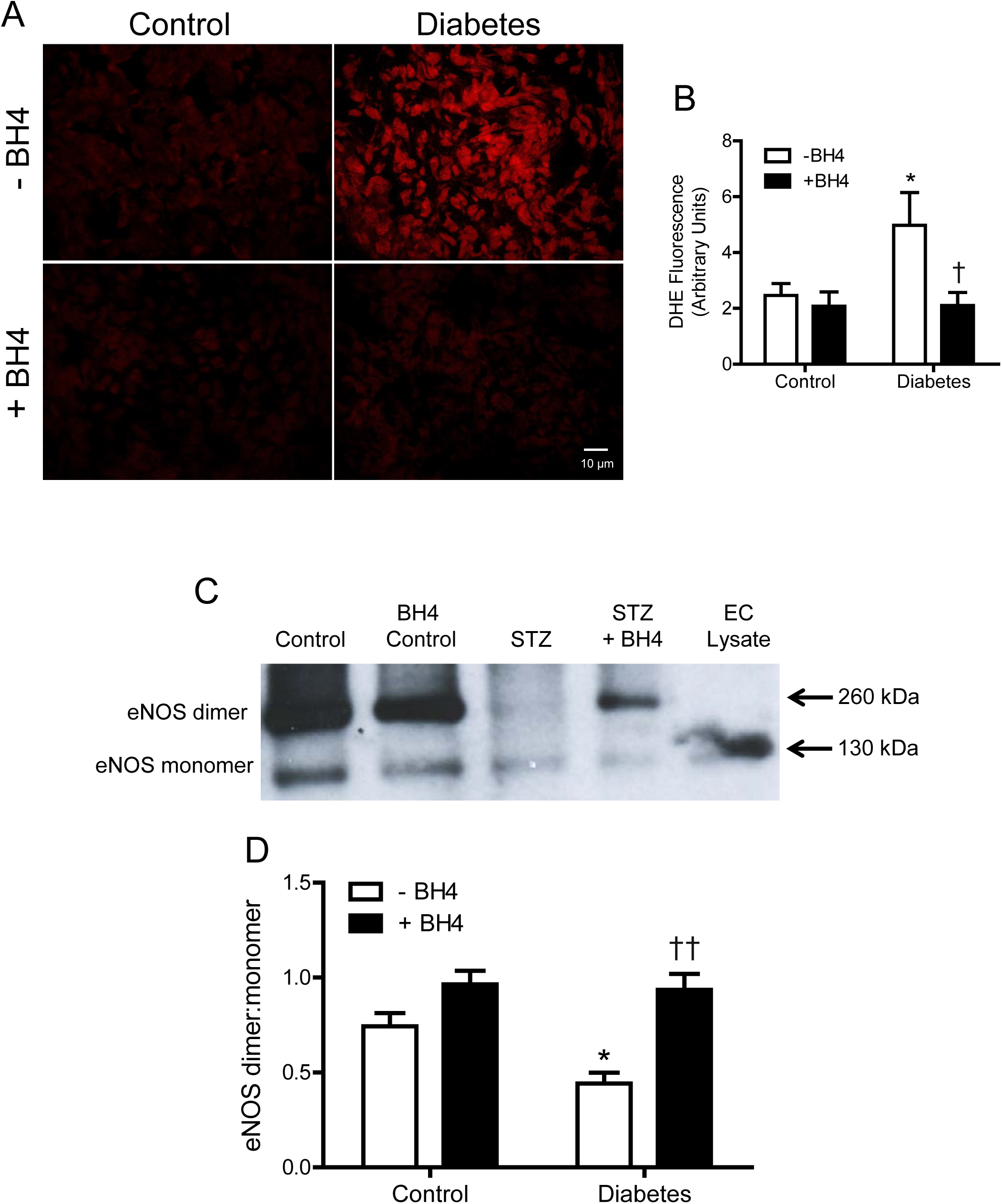
Effects of sapropterin (BH4) on superoxide production and eNOS dimer/monomer protein levels in E12.5 hearts. (A) Representative images of DHE staining in the ventricular myocardium of E12.5 hearts from control and diabetic dams with and without BH4 treatment. (B) Quantification of DHE fluorescence signals. (C) Representative Western blotting showing eNOS dimer and monomer bands of E12.5 hearts from control and diabetic dams with and without BH4 treatment. Denatured endothelial cell lysate validates the size of the eNOS monomer band. (D) Densitometric analysis of eNOS dimer/monomer ratios. Data are means ±SEM, n = 5 - 6 hearts per group. \**P*<0.05 vs. untreated control, ††*P*<0.01 vs. untreated diabetes.

## DISCUSSION

The present study utilized a clinically relevant model of CHDs induced by pregestational diabetes we recently established [27, 28]. Consistent with our previous studies, a spectrum of CHDs were observed in the offspring of diabetic mothers. The CHDs range from ASD, VSD and valve thickening, to major malformations including AVSD, DORV, truncus arteriosus and hypoplastic left heart, as clinically categorized by Hoffman *et al* [30]. Sapropterin treatment was able to prevent all major defects and significantly reduce the overall level of defects induced by pregestational diabetes. Other changes in the fetal heart such as thinner ventricular myocardium, thickened valves, stenosis of the aorta and pulmonary artery, impaired cardiac function along with decreased cell proliferation and gene expression induced by maternal diabetes, were normalized with sapropterin treatment. Furthermore, sapropterin administration in the diabetic dams increased eNOS dimerization and decreased ROS levels in the fetal heart. Our study suggests that sapropterin improves eNOS coupling and prevents CHDs in pregestational diabetes in mice (Fig. 9).

**Figure 9.**
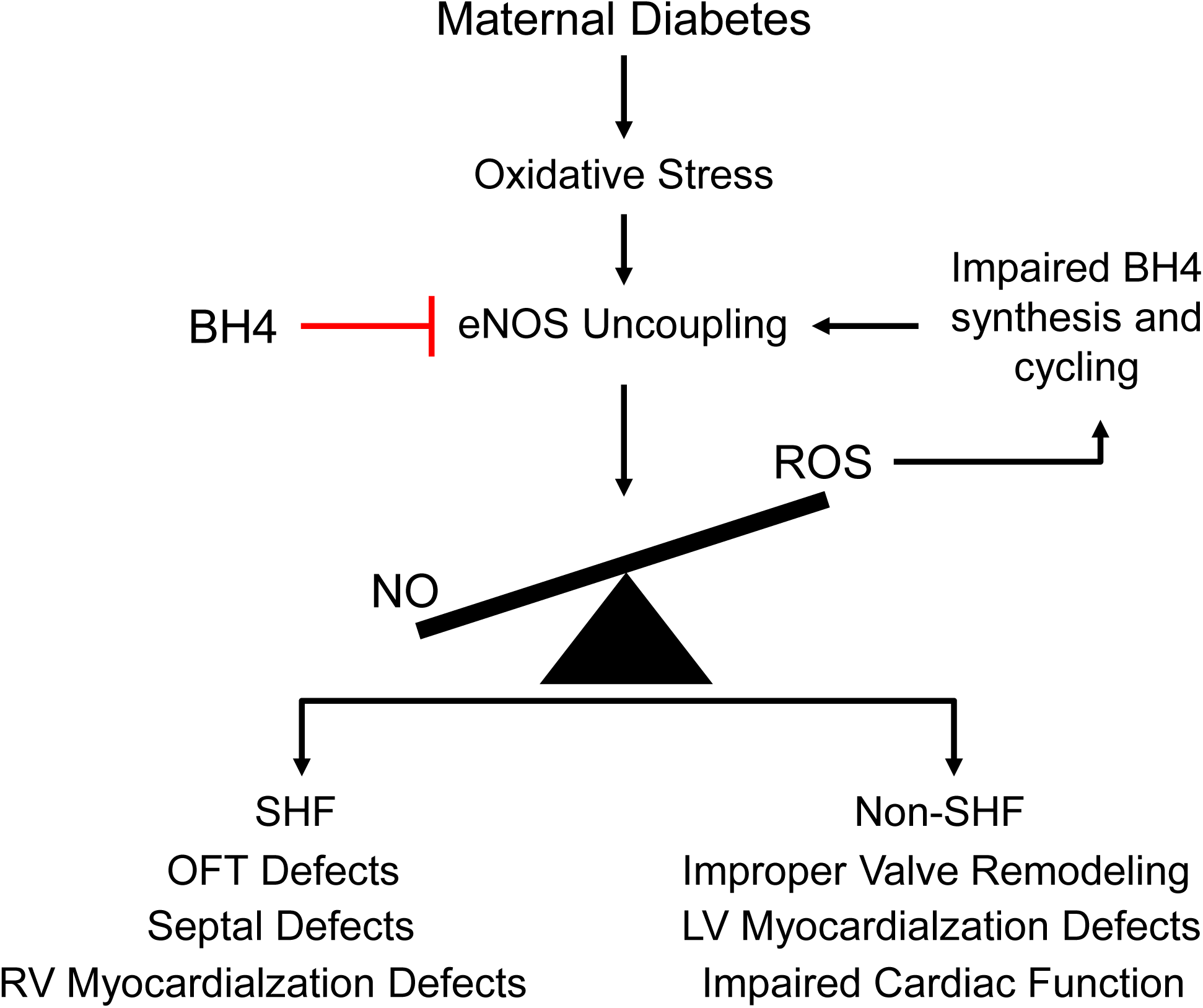
Schematic summary of eNOS uncoupling and CHDs induced by pregestational diabetes and the inhibitory effects by sapropterin (BH4) treatment. SHF, second heart field.

In this model, STZ was used to induce hyperglycemia in female mice at least 7 days before they were bred with normal healthy males. Since STZ has a short plasma half-life of 10 minutes in rodents [31], it is unlikely that STZ would cause any teratogenic effects in the fetus. To confirm that hyperglycemia is the cause of CHD in this model, a group of mice were treated with insulin to lower glucose levels in diabetic dams. Our data show that insulin treatment abrogated CHDs in pregestational diabetes, which is consistent with clinical studies that good glycemic control in women with pregestational diabetes lowers the incidence of CHDs [6]. It should be noted that the incidence of CHDs found in the diabetic groups with or without sapropterin treatment at E18.5 may be an underestimate as fetuses may have been absorbed in utero at around E12.5. To study the effects of sapropterin on maternal diabetes, blood glucose levels were assessed during pregnancy. Our data show that sapropterin treatment in the diabetic dams had no effect on the hyperglycemic state of the animal, suggesting that the beneficial effects of sapropterin on fertility (or percent successful pregnancy), litter size, and heart development under pregestational diabetes are independent of blood glucose levels. In humans, sex-related differences in the prevalence of CHDs have been reported. In a recent large cohort study with 9,727 cases of CHDs, the male to female ratio was 55% to 45%, indicating a significant predisposition of CHDs in the male sex [32]. In agreement with clinical studies, our data show a male dominance of CHDs in the offspring of diabetic mothers. Notably, sapropterin treatment appears to be more effective in preventing CHDs in females than in males.

We have recently shown that defects in SHF signaling results in thin myocardium, septal defects and OFT defects [23, 33]. The defective SHF progenitor contribution may explain the majority of heart malformations in the OFT, cardiac septum and cardiac valves induced by pregestational diabetes. Since sapropterin treatment significantly prevented these CHDs in the offspring of diabetic dams, it is likely that impairment in SHF progenitors will be diminished by maternal sapropterin treatment, which still needs to be confirmed in future studies. Considering the presence of hypoplastic left heart and OFT septation defect (truncus arteriosus), it is possible that pregestational diabetes also impairs FHF and CNC cells in our model. Notably, these defects were all prevented in diabetic dams treated with sapropterin, suggesting that sapropterin treatment improves the function of FHF and CNC progenitors in pregestational diabetes. These findings are consistent with previous studies showing that oxidative stress during diabetic pregnancy disrupts cardiac neural crest migration and causes OFT defects in rodents [34, 35], which are prevented by treatment with vitamin E, an antioxidant [36].

Cell proliferation is critical to cardiac morphogenesis and the growth of the fetal heart [37]. Decreased cell proliferation results in shortening of OFT length, thin myocardium and CHDs [33, 38]. In the present study, E10.5 hearts show less cell proliferation and a shorter OFT in embryos from diabetic dams. Furthermore, E18.5 hearts from diabetic dams are notably smaller and their ventricular walls are significantly thinner than control hearts. These abnormalities were all prevented by sapropterin treatment. Previous studies have shown that BH4 increases DNA synthesis and cell proliferation in erythroid cells [39]. Additionally, BH4 mediates the proliferative effects of epidermal growth factor and nerve growth factor in PC12 cells [40]. We have previously shown that eNOS promotes cardiomyocyte proliferation [14, 41]. Since BH4 increases eNOS coupling as shown by higher dimer/monomer ratios in our study, it is possible that a normalized eNOS signaling may contribute to the improved cell proliferation by sapropterin treatment. Increases in cell apoptosis may also contribute to CHDs. However, we have previously shown that the incidence of cell apoptosis in fetal hearts of diabetic offspring is low (about 1%) and was not affected by anti-oxidant treatment, suggesting an insignificant role of apoptosis in our model [28]. We therefore did not assess cell apoptosis in the present study.

Cardiac transcription factors including Gata4, Gata5, Nkx2.5 and Tbx5 are critical to normal heart development, and genetic mutations of these transcription factors result in CHDs in humans [42]. Interestingly, both eNOS and ROS can alter the expression of these transcription factors [13, 28]. For example, deficiency in eNOS decreases the expression of cardiac transcription factors including *Gata4* during embryonic heart development [16]. Additionally, the expression of cardiac transcription factors was downregulated in the fetal heart of offspring of diabetic mothers, which was restored by treatment with an antioxidant, N-acetylcysteine [28]. In agreement with our previous studies, we showed a downregulation of *Gata4, Gata5, Nkx2.5* and *Tbx5* in fetal hearts of offspring from diabetic mothers in the present study. Decreased expression was also seen in *Bmp10*, essential in cardiac growth and chamber maturation [43]. Importantly, sapropterin administration to diabetic dams restored the expression profile of these factors to normal levels. We also assessed the expression of *GCH1* and *DHFR*, which are enzymes responsible for de novo BH4 biosynthesis and recycling of BH2 back to BH4, respectively. Both GCH1 and DHFR are sensitive to oxidative stress and NO signaling [18, 44]. During pregestational diabetes, GCH1 and DHFR transcript levels in the fetal heart were downregulated, which was normalized by sapropterin treatment. The effects are consistent with eNOS uncoupling and ROS production induced by pregestational diabetes and the ability of sapropterin treatment to recouple eNOS and restore ROS balance within the embryonic heart.

A major non-cardiac malformation induced by pregestational diabetes is NTD such as anencephaly, exencephaly and spina bifida [45]. It has been reported that 25-40% offspring of diabetic dams have NTDs when the fetuses are examined at E10.5 [46, 47]. To analyze cardiac malformation, we examined the fetuses at E18.5 and may have missed most of the NTDs, which are likely absorbed beyond E10.5. This may be the reason that we only observed one exencephaly in the present study and a low incidence of NTDs (4.8%) in our previous study [28]. Apart from being a cofactor of eNOS, BH4 is also a substrate for aromatic amino acid hydroxylases [48]. Additionally, BH4 biosynthesis is important to neural tube development. Inhibition of GCH1 activity, a rate limiting enzyme in the biosynthesis of BH4, interrupts neural tube closure, which can be prevented by BH4 treatment in chick embryos [49]. Furthermore, *GCH1* haplotypes are significantly associated with a higher risk of NTD in infants [50]. It is possible that sapropterin treatment may prevent NTD induced by pregestational diabetes, a hypothesis that needs to be tested in future studies.

In summary, the present study demonstrates that treatment with sapropterin (Kuvan®), an orally active synthetic form of BH4 during gestation improves eNOS coupling, reduces ROS and increases cell proliferation in the embryonic heart of offspring of diabetic mothers. Notably, sapropterin treatment prevents the development of major CHDs induced by pregestational diabetes. Sapropterin is an FDA approved drug to treat phenylketonuria (PKU), a genetic disorder due to mutations of phenylalanine hydroxylase (PAH) gene leading to low levels of phenylalanine hydroxylase [22]. Our study suggests that sapropterin may also have therapeutic potential in preventing CHDs in offspring of women with pregestational diabetes.

## Acknowledgment

This study was funded in part by grants from the Canadian Institutes of Health Research (CIHR) to Q.F. and T.D., and the Children’s Health Foundation in London, Ontario, Canada to K.N., T.D. and Q.F. We thank Dr. Ben Rubin, Western University for his advice on statistical analysis. Q.F. is a Richard and Jean Ivy Chair in Molecular Toxicology at University of Western Ontario.

## Author contributions

A.E, X.L., and Q.F. conceived the experiments. A.E, X.L., B.U., T.D., K.N., and Q.F. designed the experiments. A.E., T.S., X.L. and A.K. performed the experiments and data analyses. A.E. and Q.F. wrote the paper. A.E., B.U., T.D., K.N., and Q.F. revised the manuscript. All authors contributed to the interpretation of results and proofreading of the manuscript.

